# The BRCA1-RAD51 Axis Regulates SCAI/REV3 Dependent Replication Fork Maintenance

**DOI:** 10.1101/2025.11.25.689574

**Authors:** Chloe Unterseher, Hiroshi Tsuchida, Rosevalentine Bosire, Shaylee Kieffer, Xuhui Jin, Richard O. Adeyemi

## Abstract

Replication stress is a key contributor to genomic instability and cancers. BRCA1 has well established roles in protecting stalled forks against degradation. Here we show that BRCA1 has a fork protection-independent role via regulation of RAD51 in a manner that confers dependency on SCAI and REV3. SCAI loss leads to elevated DNA breaks, genomic instability and cell death upon DNA damage. We show that the increased DNA breaks occur in S phase, consistent with fork breakage. BRCA1 drives SLX4-SLX1-ERCC1 mediated DNA break formation in the absence of SCAI and REV3, which are required to maintain integrity of stalled forks for efficient restart. Domain analysis revealed that the increased break signaling seen in the absence of SCAI does not require binding to resection factors like CtIP. Surprisingly, loss of fork reversal factors leads to additive increases in damage signaling and elevated fork shortening in the absence of SCAI/REV3 in a manner that depends on RAD51 activity. We propose that Protexin may be required to replicate through and maintain stalled replication forks at fragile genomic regions. Failure to do so leads to increased DNA breakage and genomic instability.

## INTRODUCTION

Faithful duplication of the genome of dividing cells is continually threatened by impediments that challenge replication fork progression^1^. Such obstacles can result from lesions arising from exogenous DNA damage or from endogenous sources such as DNA secondary structures, difficult-to-replicate chromatin, oncogene-driven hyper-replication, and transcription-replication conflicts, among others^2^. Collectively, these inputs result in replication stress and, if left unresolved, drive fork stalling, breakage, and collapse with subsequent genome instability—a key hallmark of several cancers^3,4^. To avert these outcomes, cells deploy a coordinated replication-stress response that stabilizes perturbed forks, remodels fork architecture, and re-establishes DNA synthesis through pathway choices that include repriming, translesion synthesis (TLS), template switching, and recombination-based restart^5^.

A central remodeling route at stalled forks is fork reversal—conversion of the stalled fork into a four-way duplex intermediate—catalyzed by annealing helicases/ATP-dependent translocases such as SMARCAL1, ZRANB3, and others^6,7^. Fork reversal can protect against further damage, yet reversed forks are intrinsically vulnerable. Homologous recombination (HR) factors, most notably RAD51 and BRCA1/BRCA2, extend their canonical roles beyond two-ended double strand break (DSB) repair to stabilize and protect nascent strands at reversed forks^8,9^. However, the varied functions of these proteins at stalled forks and how these differ from conventional two-ended breaks is still not clear.

In addition, how stalled forks transition back to productive replication remains an area of active investigation^10,11^. In addition to RAD51’s fork protective function, several studies have shown that RAD51 drives reversal of stalled forks and is important for promoting fork restart ^12,13^. Forks can restart via repriming^14^ (which suppresses fork reversal and leaves post-replicative single-stranded DNA gaps to be filled later by template switching or TLS) or via direct RECQ1 activity on reversed forks^15^. Alternatively, stalled or reversed forks can be cleaved by structure-specific endonucleases (e.g., MUS81–EME1, SLX1–SLX4), generating one-ended DSBs that are repaired via recombination-based restart^16,17^. In mammalian cells, such restart often resembles break-induced replication (BIR)—a conservative, long-tract synthesis program that, in several contexts, operates with minimal contribution from RAD51’s strand-exchange activity ^18,19^. Alternatively, RAD51 dependent recombination restart mechanisms without break formation have also been proposed^20,21^.

We previously identified a general requirement for the Protexin complex—comprising SCAI and the Pol ζ catalytic subunit REV3—in promoting viability after replicative damage^22^. In particular, we uncovered a TLS-independent function of SCAI/REV3 in the response to DNA interstrand crosslinks (ICLs), where SCAI restrains excessive EXO1-mediated resection and preserves replication competence. These findings, however, raised the possibility that SCAI/REV3 does more than modulate a single nuclease axis at ICLs but may influence other aspects stalled-fork metabolism. Here, we investigate how the SCAI/REV3 complex regulates fork advancement and how the complex interfaces with the RAD51/BRCA1 module to govern outcomes at stalled forks. We show that SCAI/REV3 is required to maintain fork integrity during replication stress and to promote efficient fork restart. In the absence of SCAI, there is increased replication stress associated DNA breakage in S phase, genomic instability and cell death. Strikingly, break formation in the absence of Protexin is orchestrated by BRCA1-RAD51 via recruitment of SLX4 to chromatin. Furthermore, combined loss of SMARCAL1 and SCAI led to additive break-induced damage signaling and defective fork maintenance. We propose that Protexin loss causes a failure of strand invasion mediated fork restart with resultant fork breakage and genomic instability.

## RESULTS

### Protexin complex prevents DNA breakage and genomic instability following replication stress

We and others previously identified SCAI as a key factor in interstrand crosslink repair^22,23^. Our previous work also suggested that SCAI might prevent cell death upon replication stress. Indeed, consistent with previous findings in U2OS cells, colony formation assays in RPE1 cells showed that SCAI depletion led to diminished viability upon treatment with hydroxyurea (HU) (Figure S1A & B). To examine how loss of SCAI impacted damage signaling following replication fork stalling, RPE1 cells were transfected with two different siRNAs to SCAI, treated with HU and blotted for replication stress markers. As expected, and consistent with what was observed following treatment with crosslinking agents^22^, SCAI loss led to increases in CHK1 and RPA2 phosphorylation (Figure 1A). Upon SCAI depletion with either siRNA, we also observed significantly increased phosphorylation of H2AX (hereafter γH2AX), typically a marker of elevated replication stress and/or DSBs, without any changes to H2AX protein amounts (Figure 1A). To further examine this, we made use of U2OS cells engineered to knock out SCAI using CRISPR. In this cell type, we observed similar increases in γH2AX and RPA2 phosphorylated on S4/8 (hereafter pRPA) (Figure 1B). Upon HU treatment, the increased pRPA in ΔSCAI cells could be rescued by re-introducing SCAI (Figure 1C, compare lanes 2, 4 & 6). To further explore SCAI’s function during replication stress, we treated cells with aphidicolin, a DNA polymerase alpha inhibitor that also causes replication stress and fork stalling^24,25^. Similar to what was observed following HU treatment, SCAI depletion led to increased γH2AX and pRPA upon treatment with aphidicolin (Figure 1D), showing that this altered signaling stemmed from a dependency on SCAI at distressed forks.

**Figure 1:**
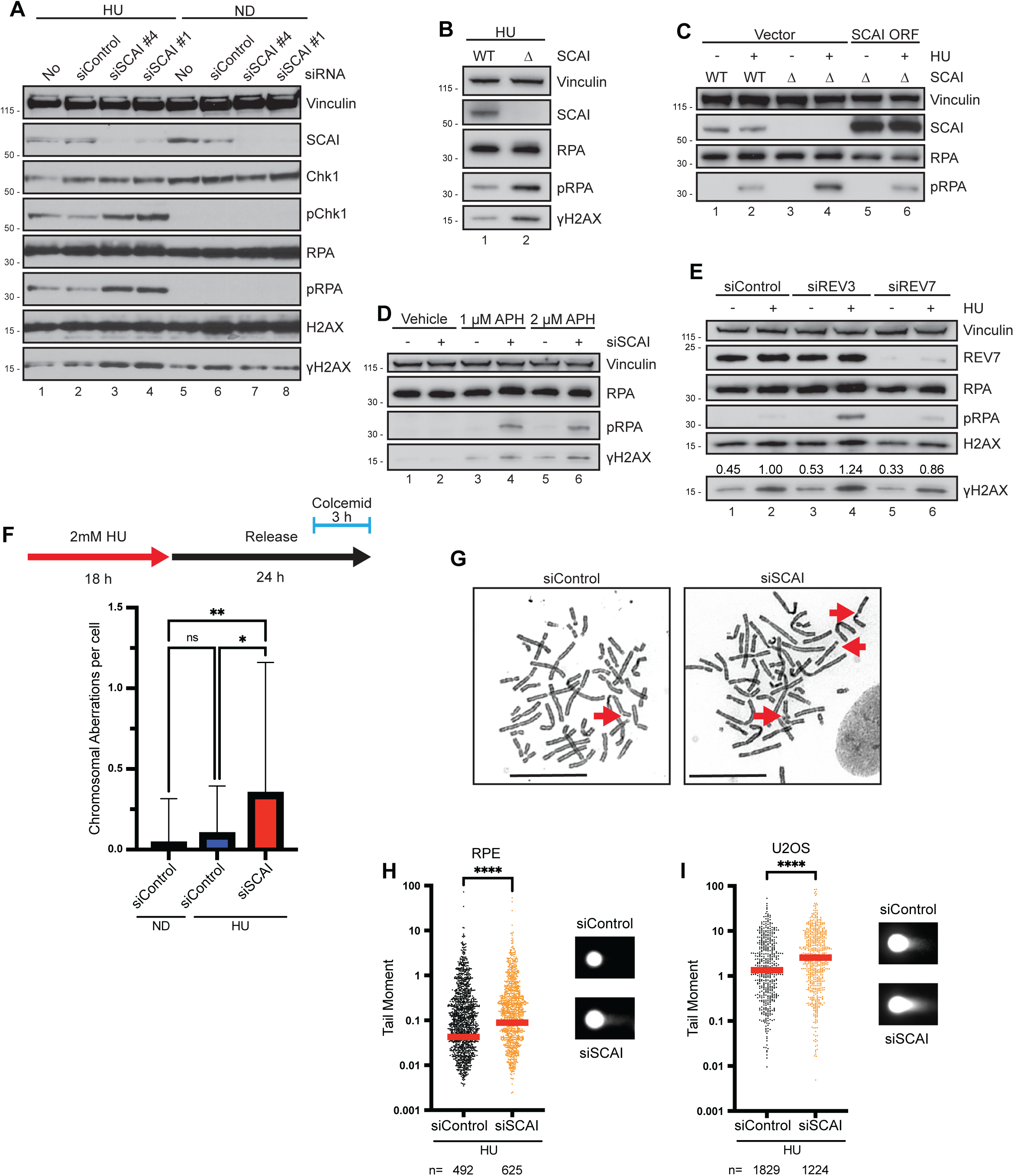
Protexin complex prevents DNA breakage and genomic instability following replication stress. **A.** WT RPE1 cells transfected with the indicated siRNAs for 60 h, then treated with 2 mM hydroxyurea (HU) for 18 h, harvested, and processed for western blotting with the specified antibodies. ND, no drug, n = 3 independent experiments. **B.** Western blots (WB) from WT and SCAI-null U2OS cell lines treated with 1 mM HU for 18 h, n = 3 independent experiments. **C.** WT and ΔSCAI U2OS cells were transfected with either backbone plasmids (Vector) or plasmids expressing SCAI that is resistant to CRISPR/Cas9 editing. After 30 h, cells were treated with 1.5 mM HU for another 18 h before immunoblotting. n = 3 independent experiments. **D.** WT RPE1 cells were transfected with the indicated siRNAs, then treated with vehicle (DMSO), 1 μM, or 2 μM aphidicolin (APH) for 18 h before being processed for western blotting with the specified antibodies. n = 3 independent experiments. **E.** WT RPE1 cells were transfected with the indicated siRNAs, then treated with 1 mM HU for 18 h and assayed for the indicated proteins by WB. γH2AX values are normalized to vinculin loading control. n = 3 independent experiments. **F.** Top: Schematic of cell treatment protocol. Bottom: Numbers of chromosomal aberrations per metaphase cell. hTERT-RPE1 cells treated with 2 mM HU for 18 h followed by washout of HU and 24 h incubation in normal medium. Indicated siRNAs were transfected two days before HU treatment. n = 3 independent experiments, mean ± S.D. Ordinary one-way ANOVA followed by Tukey’s multiple comparison test. ns – not significant (P ≥ 0.05). **G.** Representative images of (**F**). Red arrow – chromosomal aberrations. **H-I.** Comet assay of WT RPE1 (**H**) and U2OS (**I**) cells treated with 2 mM HU for 40 h. Representative comets are shown on the right. Indicated siRNAs were transfected two days before HU treatment. Red line represents median, n = 3 independent experiments, representative experiment shown. Mann-Whitney test. ****P < 0.0001.

To examine whether REV3 shared similar roles with SCAI in preventing DNA damage at distressed forks, we depleted REV3 using siRNAs. Similar to what was observed upon SCAI loss, REV3 depletion led to increases in pRPA and γH2AX upon fork stalling (Figure 1E, compare lanes 2 and 4). As an orthogonal approach, we examined replication stress signaling events using immunofluorescence (IF) microscopy. To determine whether the observed increased pRPA signified increases on chromatin, cells were pre-extracted to remove unbound proteins. Depletion of either SCAI or REV3 led to increased intensity of RPA upon HU treatment, suggesting that increased pRPA occurred on chromatin (Figure S1C). We also observed increased γH2AX using this approach (Figure S1D).

REV3 has well characterized roles in translesion synthesis^26,27^. Because HU is not thought to lead to formation of DNA adducts^28^, increased damage signaling upon REV3 loss might be independent of REV3’s role in performing TLS across damaged DNA bases. To further examine this, we depleted cells of REV7, which is critical for TLS as a component of DNA polymerase zeta^29^. Whereas REV3 loss led to increases in pRPA and γH2AX after HU treatment, REV7 depletion (and thus inhibition of TLS) did not (Figure 1E, compare lanes 4 and 6). To directly test whether failure of TLS would result in increased damage signaling we treated cells with a previously reported TLS inhibitor^30^. Similar to REV7 depletion, TLS inhibition did not increase pRPA and γH2AX after replication stress (Figure S1E) suggesting that stalled fork induced damage signaling upon REV3 loss was not attributable to a TLS function of REV3. Depletion of REV3 in SCAI knockout (KO) cells did not significantly alter the amounts of pRPA and γH2AX suggesting that both proteins function in the same pathway (Figure S1F).

The above data show that SCAI is required to prevent cell death after replication stress. The heightened signaling markers are suggestive of DNA breakage. To investigate whether SCAI loss leads to increased breaks and genomic stability after replication stress, we first treated cells with HU as before, then released from HU treatment and allowed cells to proceed into mitosis (Figure 1F, top panel). We then performed mitotic DNA spreads to examine for the presence of chromosomal abnormalities including breaks, gaps, fusions and similar events. SCAI loss did not significantly impact the extent of chromosomal abnormalities observed in the absence of HU treatment. In contrast, following treatment with HU, we observed that loss of SCAI led to significant increases in the amount of broken and otherwise abnormal chromatids compared to control knockdown cells (Figure 1F & G). The elevated abnormalities prompted us to directly examine whether SCAI limited DNA break formation. To test this, we performed neutral comet assays under a similar timeline but without releasing from HU. This should prevent mitotic entry and allow us to determine whether SCAI loss led to DNA breaks prior to mitosis. Depletion of SCAI led to significant increases in DNA break formation following replication stress compared to controls in both RPE1 (Figure 1H) and U2OS cells (Figure 1I). Similar results were seen with REV3 depletion (shown later in Figure 4E). Taken together, these data show that the Protexin complex is required for efficient repair and viability following replication stress and that its loss results in elevated DNA breakage and heightened genomic instability.

### SCAI/REV3 prevent DNA breakage in S phase

Having shown that SCAI loss upon replication stress led to increased damage signaling with increased DNA breakage, we next sought to understand the kinetics and determine when the DNA breaks form. We hypothesized that the breaks resulted from distressed forks and thus occurred in S phase. We first performed IF assays using well established fixation protocols that allow detection of PCNA specifically in S phase^31^. These experiments were done over a time course (Figure S2A-C). We co-stained for γH2AX and, as expected, observed increased γH2AX in SCAI KD cells by 16 following treatment with HU compared to controls (Figure S2A-C). There was high concordance between cells in S phase and cells exhibiting γH2AX staining (Figure S2D-E). At 8 h following HU treatment, only around 25% of cells on average were PCNA-positive, however, when comparing the observed frequency of PCNA/γH2AX double-positive cells to the expected overlap based on independent staining frequencies, we detected a significant enrichment of co-staining that was more pronounced upon SCAI depletion (Figure 2A & B). These findings indicate that the increased DNA damage signaling observed upon HU treatment in SCAI KD cells occurs predominantly in S-phase. Performing similar experiments by immunoblotting, pChk1 was rapidly seen 4h after HU treatment (Figure S2F). pRPA and γH2AX did not however appear until later (Figure S2F). Because western blotting examines the entire population of cells, many of which are outside of S phase, we synchronized cells at the G1/S boundary using double thymidine block and performed a similar time course. Under these conditions, loss of SCAI rapidly led to significant increases in γH2AX by 4 - 8h after HU treatment (Figure 2C). Parallel cell cycle analyses revealed that these cells were largely in S phase at this time (Figure S2G) and would remain so due to HU-induced S phase arrest^32^.

**Figure 2:**
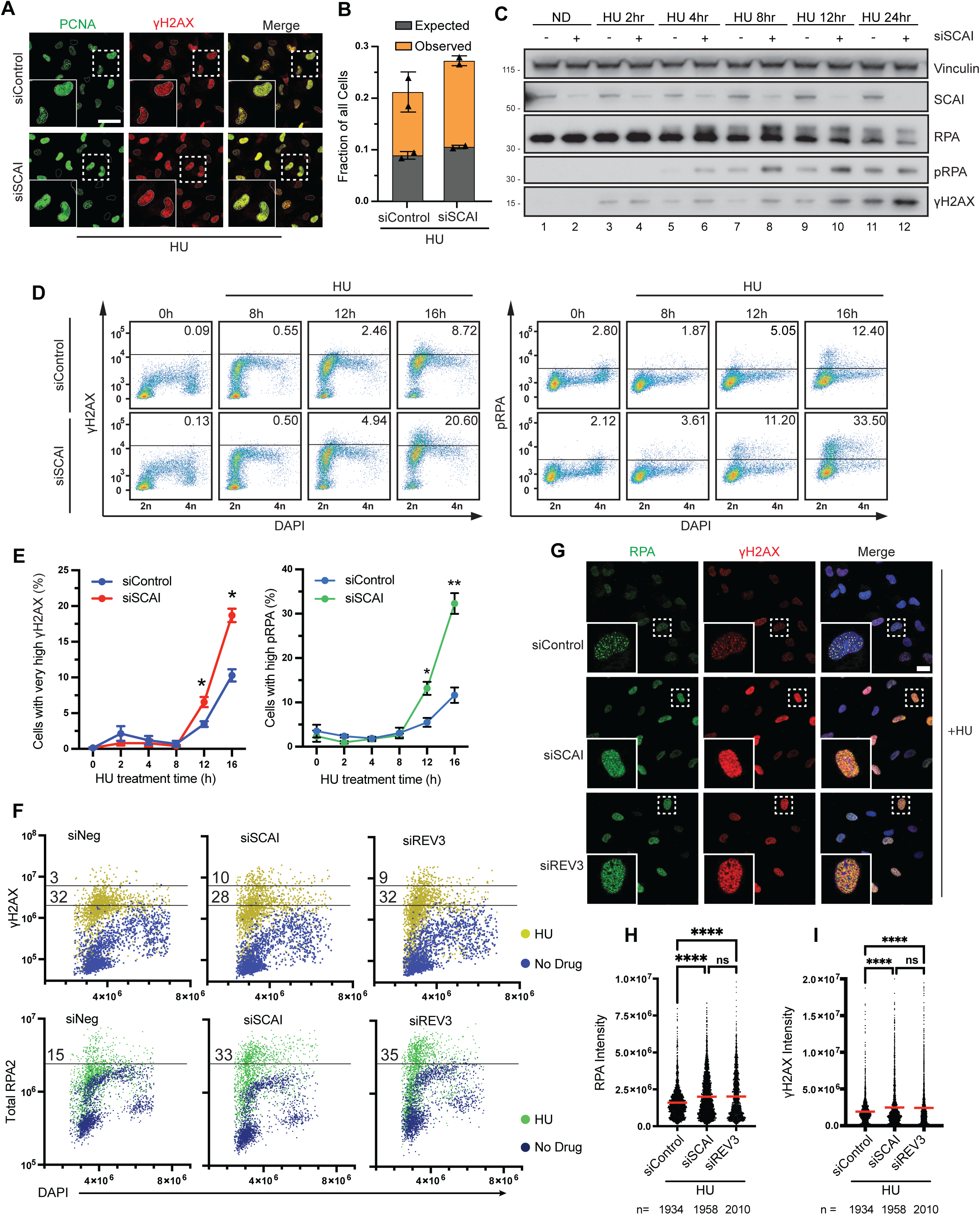
SCAI/REV3 prevent DNA breakage in S phase. A. Representative IF images showing increased γH2AX in PCNA positive compared to PCNA negative RPE cells 8 h after treatment with 1.5 mM HU. Cells were transfected with control or SCAI siRNA 2 days prior. Scale bars indicate 50 micron. B. Plot showing the observed overlap between γH2AX and PCNA staining from (A), n = 2 independent experiments, with >180 cells/condition per experiment. Expected values were obtained by multiplying γH2AX and PCNA positive fractions. **C.** RPE1 cells were synchronized by double thymidine block, released for 2 h and treated with 2 mM HU for the time points shown prior to immunoblotting. Representative experiment shown, n = 2 independent experiments. **D.** Representative scatter plots showing time course changes in cell cycle distribution and level of γH2AX (left panel) or pRPA (right panel) in RPE1 cells transfected with control or siRNA targeting SCAI then treated with 1.5 mM HU for the indicated duration. The percentage of cells with high pRPA and γH2AX signals is indicated in each panel. **E.** Quantification (mean ± SEM) of three experiments as in (**D**), unpaired t-test, n = 3 independent experiments. *P=0.0332, **P = 0.002. **F.** Multivariate scatter plots from QIBC analysis showing changes in cell cycle distribution and level of γH2AX (top panel) and chromatin-bound RPA2 (bottom panel) in RPE1 cells transfected with either control, or siRNA targeting SCAI or REV3 then treated with 1.5 mM HU for 16h. Top panel: untreated control (light blue), HU treated (yellow). Bottom panel: Untreated control (navy blue), HU treated (green). The numbers on the plots show the percentage of HU treated cells whose γH2AX or chromatin bound RPA2 was above the baseline in the no drug controls. **G**. Representative microscopy images showing increased chromatin bound RPA (green) and γH2AX (red) upon Protexin loss. Scale bar indicates 25 micron. **H-I**. Quantification of the integrated fluorescent intensity of RPA (**H**) and γH2AX (**I**). Red bar indicates mean and SEM of about 2000 nuclei. ****P < 0.0001, ANOVA followed by Tukey’s.

To directly determine whether the increased signaling occurred in S phase populations, we performed flow cytometry experiments where we coupled cell cycle analysis using DAPI with staining for γH2AX and pRPA over a 16-hour time course. This analysis showed a negligible population of cells with increased γH2AX at baseline. These mostly had near 4N DNA content indicating that these cells were either in G2 or M phase. By 8h following treatment with HU however, we observed rapid increase in cells exhibiting phosphorylated H2AX in both WT and SCAI knockdown cells (Figure 2D). Based on their DNA content, these increased population appeared to be of cells in S phase. Over the course of treatment, while the absolute numbers of cells exhibiting γH2AX staining were not significantly different at early time points, by 12 after treatment onwards, SCAI knockdown cells exhibited significant increases in the population of cells showing γH2AX staining (Figure 2D-E). Importantly, a population of cells exhibiting abnormally high levels of γH2AX was further increased upon loss of SCAI (Figure 2D-E). pRPA S4/8 is sometimes used as a marker for DNA breakage levels^33,34^ and, examining pRPA, both by western and flow cytometry, we observed similar changes in pRPA that mirrored γH2AX accumulation (Figure 2C-2E).

To further understand how DNA damage signaling was impacting cells in S phase upon SCAI or REV3 loss, we performed quantitative image-based cytometry (QIBC)^34^. This was done following pre-extraction of unbound proteins to examine chromatin bound RPA and γH2AX. We found that loss of either SCAI or REV3 led to increased amounts of chromatin associated RPA (Figure 2F-H) and, consistent with the IF data, increased population of γH2AX that was occurring in cells with S phase DNA content (Figure 2F-H). Compared to flow cytometry, QIBC affords the added advantage of visually comparing intensities of signaling on a per cell basis. Loss of either SCAI or REV3 led to increased intensity of γH2AX and chromatin associated RPA by 16 h after HU treatment (Figure 2G-H). To determine whether the increased γH2AX staining was occurring in the proximity of active replication forks, we performed proximity ligation assays (PLA) using antibodies against PCNA and γH2AX. For this assay we employed our optimized a protocol wherein PCNA is only detected in S phase. In the absence of HU, we failed to observe substantial association between PCNA and γH2AX (Figure S2H-I). Upon HU treatment however, we observed strong increases in PLA signals indicating close association between PCNA (and thus replication forks) and γH2AX staining (Figure S2H-I). PLA did not show differences between SCAI and control KD cells despite increased γH2AX staining seen by several other approaches (Figure 2). This was not surprising since HU generally causes moderate γH2AX (see Figure 2D) and PLA includes an amplification step that could minimize differences. Taken together, these experiments in which we observed increased chromatin bound total and pRPA as well as increased γH2AX in S phase cells upon Protexin loss, are suggestive of increased fork breakage. To directly examine whether SCAI loss induced increases in S-phase DNA breakage, we performed neutral comet assays following a similar cell synchronization protocol as above (Figure S2G). We found that loss of SCAI led to increases in DNA breakage 24 h after HU treatment (Figure S2J), when the vast majority of cells where still in S phase. Taken together, using a combination of approaches, we demonstrate that Protexin loss leads to increased DNA breakage in S phase.

### Protexin complex is required to maintain fork integrity during replication stress

Our observed increases in S phase DNA damage and break formation suggests a dependency on Protexin to maintain fork integrity. To directly investigate SCAI’s role in replication fork dynamics, we performed DNA fiber analysis. We pretreated cells with the first label, added HU for similar duration as our previous experiments, then released from HU while changing labels for a short duration (Figure 3A, top). This setup allows us to measure different potential effects of SCAI loss following replication stress. Under this setup, SCAI loss led to reduced lengths of both the first and second labels suggesting that SCAI was required for maintaining fork integrity upon replication stress (Figure 3B). Knockdown of REV3 also led to a similar phenotype (Figure S3A). Reduced length of the second label suggested that SCAI influenced the efficiency of fork restart. Thus, we quantified the percentage of restarted (dual labeled) forks. We observed modest reduction in the frequency of restart events upon SCAI loss (Figure 3C). SCAI’s role appeared to be limited to ongoing distressed forks, as SCAI depletion did not affect the lengths (Figure S3B, left panel) or frequency (Figure S3B, right panel) of new origin firing events (fibers incorporating only the second label) after HU release. Our observation that SCAI/REV3 loss caused reduced lengths of the first label under prolonged replication stress (Figure 1A, left panel) suggested that fork advancement during or following conditions of replication stress was compromised by loss of Protexin. To further understand SCAI’s role under these conditions, we switched our labeling strategy to introduce the second label along with HU treatment. SCAI loss consistently led to reduced fork lengths upon replicative stress induction (Figure 3D, right panel) with no significant effect on the first label preceding HU treatment and fork stalling (Figure 3D, left panel). Taken together, these data show that SCAI is required to maintain stalled fork integrity under and to promote efficient restart following prolonged replication stress.

**Figure 3:**
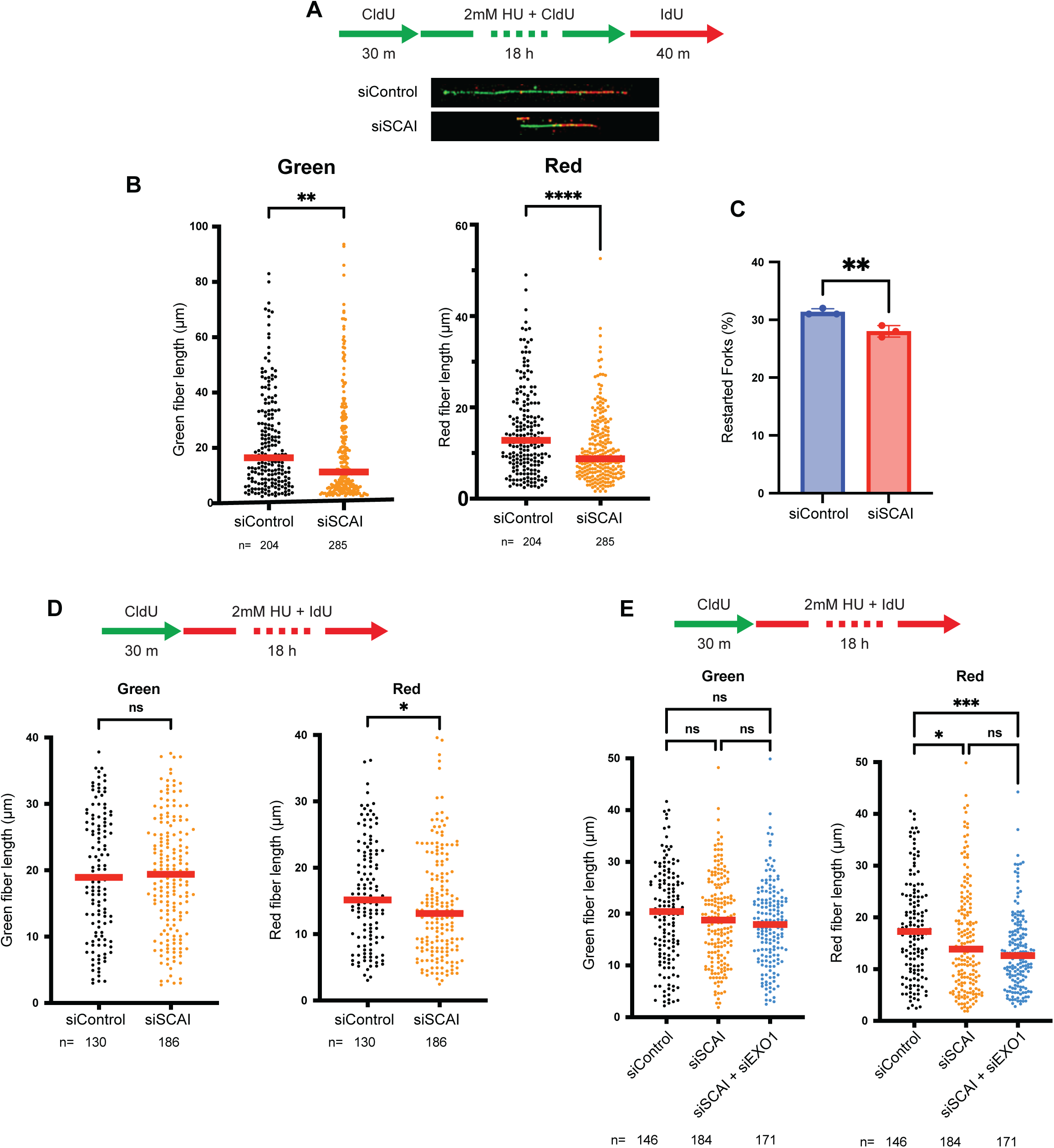
Protexin complex is required to maintain fork integrity during replication stress. **A.** Schematic of pulse-labeling DNA strands and representative images of two-color DNA fibers. DNA fibers were from RPE1 cells transfected with siRNAs as indicated. **B.** Lengths of two-color DNA fibers from (**A**). Green portion quantified in left panel and red (restarted) portion on right. **C.** Percentage of restarted forks (dual-labeled) over total amounts of green labeled forks from experiment in **A**, n = 3 independent experiments, unpaired t-test, **P = 0.0075 **D.** Top, Schematic of pulse-labeling DNA strands. **B, D-E**. Red line represents median, three independent experiments, representative experiment shown. Mann Whitney test for two group comparison and Kruskal-Wallis followed by Dunn’s multiple comparison test for more than three group comparison. ns – not significant, *P < 0.05, **P < 0.01, ***P < 0.001, ****P < 0.0001.

It is known that absolute blocks to fork advancement can lead to fork reversal with subsequent fork degradation^7,35,36^. We and others have previously reported that SCAI regulates EXO1 activity^22,37^. To examine this further, we co-depleted EXO1 and SCAI and performed similar fork dynamics assays as in Figure 3D. As expected, SCAI loss led to reduced fork lengths upon replication stress (Figure 3E, right panel) without affecting lengths under untreated conditions (Figure 3E, left panel). Surprisingly, co-depletion of EXO1 had no effect on fork lengths in the absence of SCAI under these conditions (Figure 3E), suggesting that the fork defects could not be fully explained by EXO1-mediated fork degradation. Perhaps consistent with this finding, EXO1 depletion failed to rescue HU treatment-mediated increases in pRPA seen upon SCAI loss (Figure S3C). Overall, these results demonstrate that Protexin was required for efficient fork advancement upon and following replication stress. This function of SCAI appeared to not be strongly impacted by loss of EXO1.

### BRCA1 promotes S-phase DNA break formation in the absence of Protexin

BRCA1 has multiple roles in regulating stalled fork dynamics^38,39^. Thus, we investigated whether BRCA1 might influence the increases in pRPA and γH2AX observed in the SCAI knockdown cells. To test this, we made use of RPE1 cell lines in which BRCA1 had been eliminated using CRISPR and examined for alteration in pRPA and γH2AX levels. As expected, SCAI depletion led to increased damage signaling upon HU treatment (Figure 4A, compare lanes 2 & 3) in BRCA1 proficient RPE1 cells. Strikingly, loss of BRCA1 abolished the increased γH2AX and pRPA seen upon SCAI loss (Figure 4A, compare lanes 3 & 5). As an alternative approach, we depleted BRCA1 using siRNA. Consistent with our KO data, the increased pRPA and γH2AX seen upon SCAI loss was eliminated by BRCA1 depletion (Figure S4A). BARD1 is a key interaction partner that forms a stable complex with BRCA1^40^. We depleted cells of BARD1 and, as expected, observed diminished levels of BRCA1 (Figure S4A). Basal levels of pRPA and γH2AX were higher in BARD1 knockdown vs BRCA1 knockdown cells for unclear reasons, nevertheless, BARD1 knockdown dampened increases in pRPA and γH2AX upon SCAI loss (Figure S4A). Depletion of either BRCA1 or BARD1 also eliminated increased pRPA upon cisplatin treatment (Figure S4B). It is worth noting that BRCA1 loss did not cause similar reductions in pCHK1 activation (Figure 4B and Figure S4B) suggesting that ATR activity *per se* (which regulates damage signaling upon replication stress) was not affected by loss of BRCA1.

**Figure 4:**
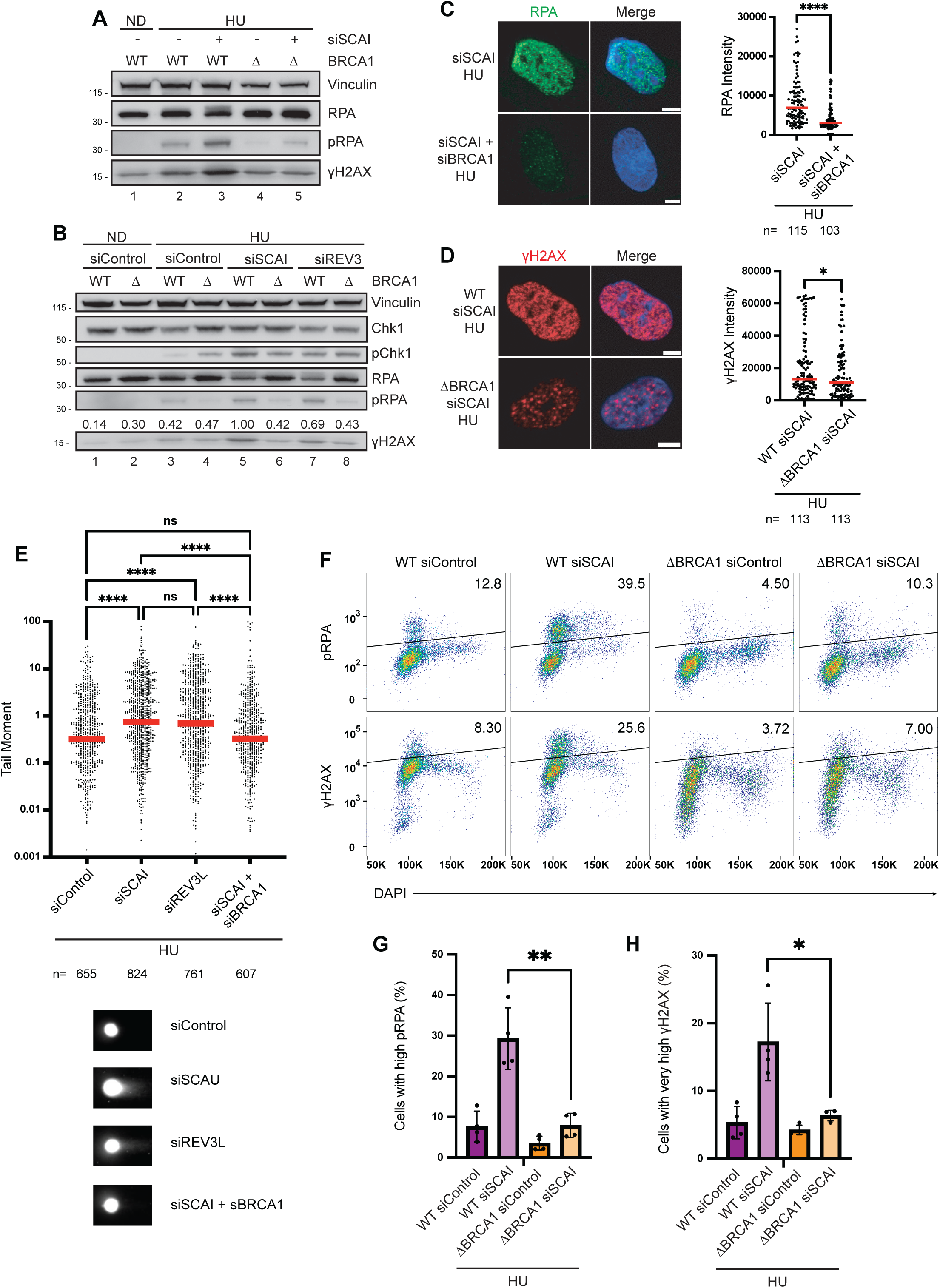
BRCA1 promotes S-phase DNA break formation upon Protexin loss. **A.** WT and BRCA1 null RPE1 cells were transfected with the indicated siRNAs, then treated with 1 mM HU for 18 h prior to blotting for the indicated proteins. Representative experiment shown, n = 3 independent experiments. **B.** WT and BRCA1-null RPE1 cells were transfected with the indicated siRNAs. Cells were then treated with 1 mM HU for 18 h before immunoblotting against the indicated proteins. Representative experiment shown, n = 3 independent experiments. **C-D.** Representative IF images showing RPA foci (**C**) and γH2AX foci (**D**) in WT RPE1 cells following treatment with the indicated siRNAs. Cells were treated with 1.5 mM HU for 18 h prior to IF. Scale bars indicate 5 micron. Quantification shown to the right. Red line represents median. **** P < 0.0001, *P = 0.0337, unpaired Welch’s t test (two-tailed). **E.** Comet assay of RPE1 cells treated with 2 mM HU for 40 h. Representative comets are shown on the bottom. Indicated siRNAs were transfected two days before HU treatment. Red line represents median, n = 2 independent experiments, representative experiment shown. Kruskal-Wallis followed by Dunn’s multiple comparison test. ****P < 0.0001, ns – not significant. **F-H.** Representative flow cytometry of WT and BRCA1-null RPE1 cells treated with 1.5 mM HU for 16 h. Indicated siRNAs were transfected two days before HU treatment. Percentage of cells with high pRPA and γH2AX signals is indicated in each panel. Quantification of percentage of cells with high pRPA (**G**) and γH2AX (**H**), n = 4 independent experiments. **P = 0.0067, *P = 0.0307, unpaired Welch’s t test (two-tailed).

To see if a similar effect applied to REV3, we depleted SCAI and REV3 using siRNAs in WT or BRCA1 KO cells. As shown earlier, we observed increases in damage signaling markers indicative of DNA breakage and resection upon loss of either SCAI or REV3 (Figure 4B, compare lane 3 with lanes 5 & 7). Importantly, cell lines lacking BRCA1 failed to show such increases (Figure 4B, compare 5 & 6 and lanes 7 & 8), suggesting that BRCA1 mediates heightened damage signaling in the absence of the Protexin complex. These findings were further confirmed using IF experiments, where loss of BRCA1 led to reduced amounts of RPA on chromatin as well as decreased γH2AX levels, again demonstrating a key role for BRCA1 in mediating damage signaling upon Protexin loss (Figure 4C-D). We showed earlier that SCAI loss leads to elevated DNA breakage (Figure 1J-K). To directly examine if BRCA1 regulates DNA breakage after SCAI/REV3 depletion, we performed neutral comet assays. As expected, loss of either SCAI or REV3 led to increased accumulation of DNA breaks (Figure 4E). Importantly, co-depletion of BRCA1 alongside SCAI prevented DNA break accumulation (Figure 4E), suggesting that the DNA breakage seen in the absence of SCAI was dependent on BRCA1 activity.

SCAI binds to ssDNA^23^. We were curious whether BRCA1 loss influenced SCAI’s recruitment to damaged chromatin. We treated WT or BRCA1 deficient cells with HU and performed chromatin fractionation experiments. As expected, absence of SCAI caused increased γH2AX (Figure S4C, compare lanes 2 & 3), that was BRCA1 dependent (Figure S4C, compare lanes 3 & 5). Importantly, SCAI accumulated in chromatin fractions upon HU treatment (Figure S4C, compare lanes 1 and 2) but this increase in SCAI accumulation was not impacted by BRCA1 loss (Figure S4C, compare lanes 2 & 4).

Finally, to directly examine whether BRCA1 orchestrated DNA breakage in S phase, we performed flow cytometry experiments. We tracked DNA content using DAPI and stained for pRPA and γH2AX. As shown earlier, in WT cells, SCAI depletion led to increases in both pRPA and γH2AX in S-phase populations (Figure 4F-H). Strikingly, this increase was abolished upon BRCA1 depletion (Figure 4F-H). Comet assays performed on synchronized S phase populations also showed DNA breaks induced upon SCAI loss was abolished following BRCA1 depletion (Figure S4D). Importantly, the BRCA1 effect was not due to cell cycle dysregulation, as BRCA1 loss by itself did not prevent entry into S phase (Figure S4E) except for minor increases in G2 populations as has been reported previously^41,42^. Taken together, our results demonstrate that BRCA1 promotes DNA breakage in the absence of Protexin following induction of replication stress.

### Defining BRCA1 requirements for regulating stalled fork breakage

BRCA1 forms several complexes important for resection control, HR and other roles; these different complexes often mediate the various functions of BRCA1^43,44^. For example, the BRCA1-CtIP complex, formed via CtIP binding to the C-terminus of BRCA1, controls resection at ionizing radiation-induced DSBs^45,46^. Since BRCA1 controls DNA breakage upon SCAI loss, we wanted to determine which subcomplexes and/or function of BRCA1 mediated this role. We first depleted ABRA1, BRIP1 or CtIP (key members of the BRCA1-A, B or C complexes) respectively. Unlike BRCA1 depletion, depletion of none of these factors prevented increased pRPA in the absence of SCAI (Figure 5A). We did however observe that depletion of Abraxas by itself increased the basal level of pRPA under HU treatment conditions (Figure 5A, lane 4). Such a role for ABRA1 (Abraxas) has been reported following treatment with the topoisomerase inhibitor, camptothecin^47^. Similar results were seen upon cisplatin treatment (Figure 5B).

**Figure 5:**
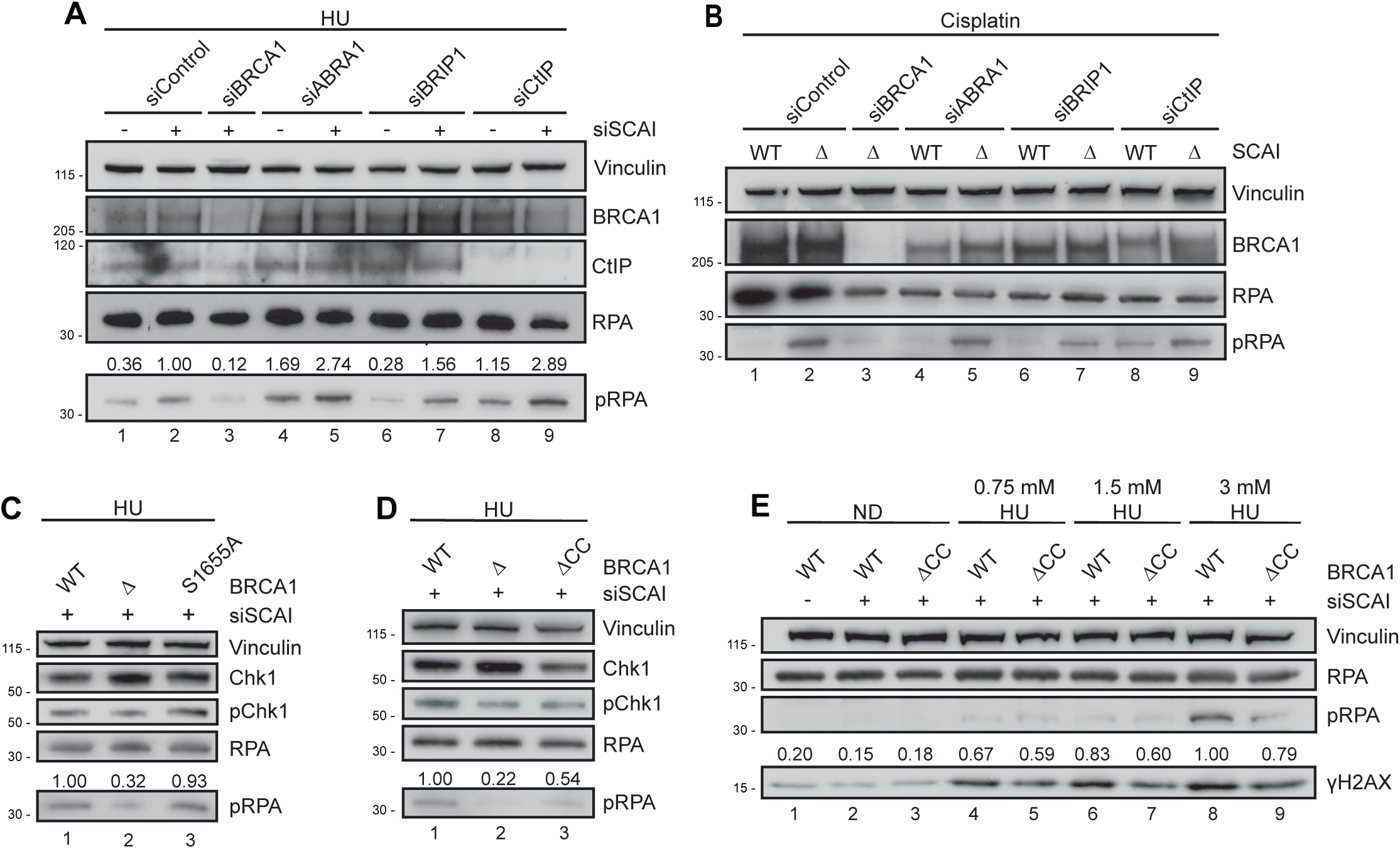
BRCA1 binding to CtIP is not required for damage signaling upon SCAI loss. **A.** WT RPE1 cells were transfected with the indicated siRNAs, then treated with 1 mM HU for 18 h and assayed for the indicated proteins by WB. Representative experiment shown, n = 3 independent experiments. **B.** Western blots from WT and SCAI-null U2OS cell lines transfected with the indicated siRNAs and then treated with 1.5 μM cisplatin for 18 h. Representative experiment shown, n = 3 independent experiments. **C.** GFP-WT-BRCA1 rescue (labeled WT), BRCA1-null (Δ), and GFP-BRCA1-S1655 (S1655) RPE1 cells were transfected with the indicated siRNAs, then treated with 2 mM HU for 18 h. WCLs were assayed for the indicated proteins by WB. Representative experiment shown, n = 3 independent experiments. **D.** GFP-WT-BRCA1 rescue (WT), BRCA1-null (Δ), and GFP-BRCA1-ΔCC (ΔCC) RPE1 cells were transfected with the indicated siRNAs, then treated with 2 mM HU for 18 h and assayed for the indicated proteins by WB. Representative experiment shown, n = 3 independent experiments. **E.** Western blots from GFP-WT-BRCA1 rescue (WT), and GFP-BRCA1-ΔCC (ΔCC) RPE1 cell lines transfected with the indicated siRNAs and treated with ND, 0.75 mM, 1.5 mM, or 3 mM HU for 18 h. Representative experiment shown, n = 3 independent experiments.

Next, we examined domain requirements of BRCA1 for DNA breakage upon SCAI loss using previously reported BRCA1 mutant-expressing cell lines^48^. Increased BRCA1 dependent damage signaling upon loss of either SCAI or REV3 could be almost fully rescued by reintroducing tagged WT-BRCA1 into knockout cell lines (Figure S5A-B). We then examined the role of two domain mutants of BRCA1. First, a point mutation (S1655A) in the BRCT domain of BRCA1 that abrogates its interaction with phosphorylation-dependent binding partners (ABRA1, BRIP1 or CtIP). A second mutant deleted the C-terminal coiled coil domain that is essential for HR. As expected, BRCA1 deletion reduced pRPA levels (Figure 5C, lane 2). Reintroduction of the BRCA1^S1665A^ mutant allowed restoration of pRPA to almost the same extent as seen following reintroduction of WT-BRCA1 (Figure 5C, compare lanes 1 and 3), suggesting that this domain was not essential for the BRCA1 induced DNA damage phenotype. This finding is consistent with what was observed following siRNA depletion of ABRA1, BRIP1 or CtIP, whose depletion failed to abrogate BRCA1’s role in promoting fork resection in the absence of SCAI (Figures 5A-B). In contrast, the coiled coil domain mutant (ΔCC) of BRCA1 did not fully restore pRPA (Figure 5D, compare lanes 1 & 3) suggesting that this domain was important for BRCA1’s role. To address this fully, we performed a dose curve of HU treatment and compared the ability of the WT BRCA1 and BRCA1^ΔCC^ mutants to rescue SCAI dependent DNA breakage and resection (Figure 5E). These data showed that the coiled coil domain of BRCA1 was required to fully orchestrate damage signaling in the absence of SCAI (Figure 5E, compare lanes 4&5, 6&7, 8&9) and that interaction between BRCA1 and CtIP does not account for the observed SCAI-depletion phenotype.

### Preventing fork reversal upon fork stalling confers a dependency on Protexin

Our data so far suggests that BRCA1 drives DNA breakage at stalled forks upon replication stress. BRCA1 also has a well-established role in protecting arrested forks from degradation^49^. This difference in phenotypes may result from differential processing by remodeling enzymes. The fork reversal activity of the enzymes SMARCAL1, ZRANB3 and HLTF represent a major mechanism of stalled fork processing^6,7^. Multiple studies have shown that when fork progression is halted, fork reversal by these enzymes drives the creation of a chicken-foot structure that is protected from breakage and resection by BRCA1^35,36^, a phenotype opposite to the DNA break promoting activity of BRCA1 that we see under our conditions. We reasoned that if DNA breakage seen in the absence of SCAI was occurring downstream of fork reversal, it would be abolished upon depletion of fork reversal enzymes. For these experiments, we have employed pRPA and γH2AX as a readout.

To determine whether fork reversal plays a similar role under our treatment conditions, we co-depleted SCAI and either SMARCAL1 or ZRANB3. To our surprise, rather than reduce damage signaling upon SCAI loss, knockdown of either SMARCAL1 or ZRANB3 led to consistent increases in pRPA and γH2AX upon HU treatment (Figure 6A, compare lanes 2, 5 & 8). The same phenotype was also seen upon co-depletion of REV3 and either SMARCAL1 or ZRANB3 (Figure 6A, compare lanes 3, 6 & 9). These results were further validated by knocking down SMARCAL1 with two independent siRNAs, which led to similar increases in the absence of SCAI (Figure 6B, compare lanes 2, 4 & 6). That this phenotype was shared between SMARCAL1 and ZRANB3 and was seen with multiple siRNAs against SMARCAL1 suggested that not only was damage signaling and presumably DNA breakage in the absence of SCAI not occurring downstream of fork reversal, but that it might also compete with fork reversal for repair of stalled forks. An alternative explanation might be that DNA breaks form downstream of a parallel fork reversal mechanism to that orchestrated by SMARCAL1/ZRANB3. Such a parallel mechanism has been suggested to occur via FBH1^50^. To examine this, we knocked down FBH1 using siRNAs. We observed that, similar to loss of SMARCAL1, FBH1 depletion also led to increased pRPA and γH2AX in the absence of SCAI (Figure 6C). Thus, depletion of multiple fork reversal enzymes all led to increased damage signaling in the absence of Protexin. That the increased damage signaling in the absence of fork reversal was more marked in the absence of Protexin suggests that a SCAI/REV3-dependent process might compete with fork reversal as a mechanism for stalled fork rescue. We cannot however rule out the possibility that Protexin and fork reversal may act at different steps of the repair process, leading to more severe phenotypes upon co-depletion.

**Figure 6:**
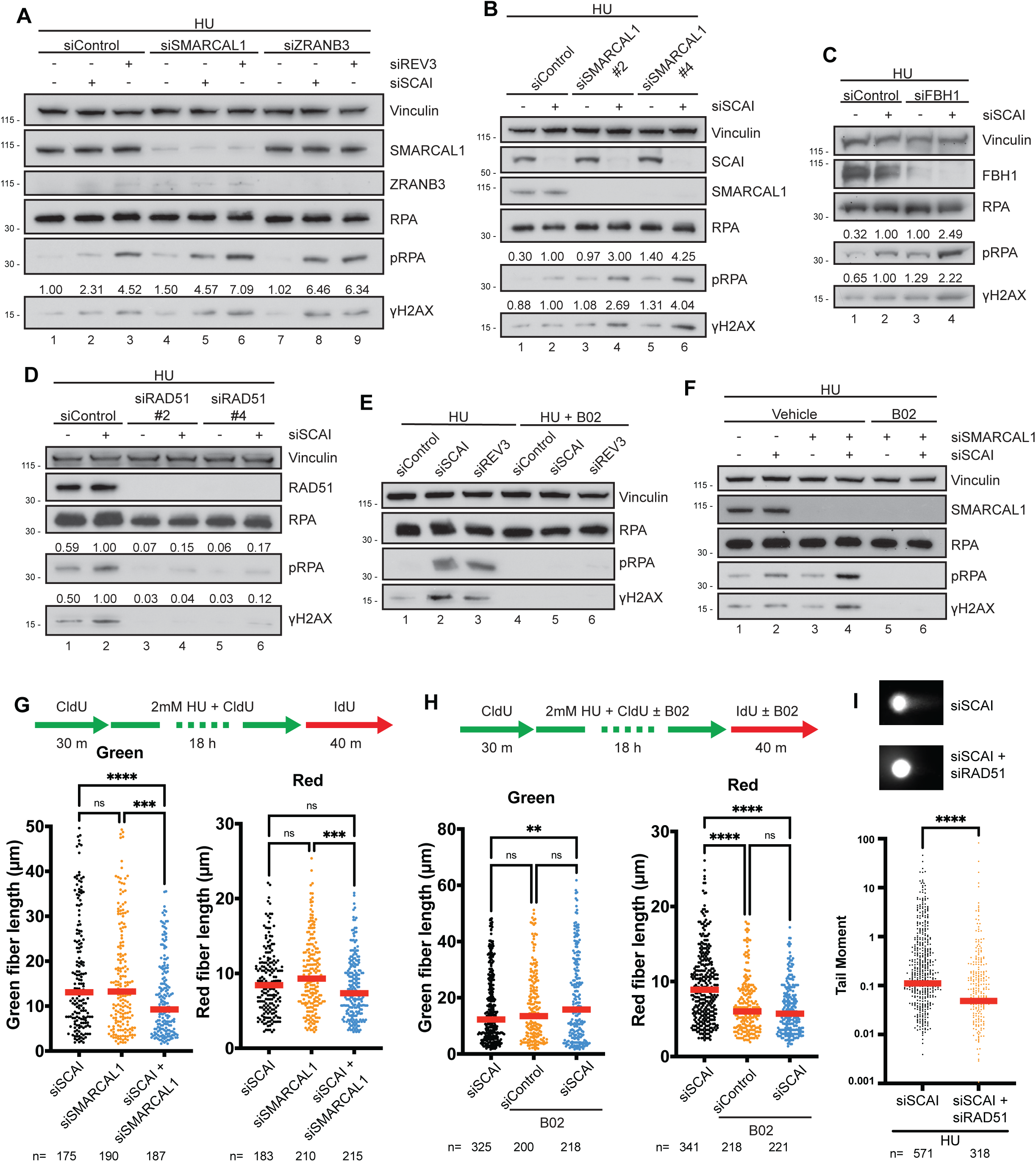
Both Protexin and reversal factors limit breaks upon replication stress. **A.** Western blots from WT RPE1 cells transfected with the indicated siRNAs and then treated with 1 mM HU for 18 h, n = 3 independent experiments. **B.** WT RPE1 cells were transfected with the indicated siRNAs, then treated with 1 mM HU for 18 h and assayed for the indicated proteins by WB, n = 2 independent experiments. **C.** WT RPE1 cells were transfected for and treated with 1 mM HU for 18 h. WCLs were prepared, and WB were performed against the indicated proteins, n = 3 independent experiments. **D.** Western blots from WT RPE1 cells transfected with the indicated siRNAs and then treated with 2 mM HU for 18 h, n = 2 independent experiments. **E.** Western blots against the indicated proteins from WT RPE1 cells transfected, then pretreated with 27 μM RAD51 inhibitor B02 or vehicle (DMSO) for 30 m and treated with 1 mM HU for 18 h, n = 3 independent experiments. **F.** Western blots from WT RPE1 cells transfected with the indicated siRNAs, pretreated with 27 μM RAD51 inhibitor B02 or vehicle (DMSO) for 30 min, then treated with 1 mM HU for 18 h. n = 3 independent experiments. **G-H**. Top: schematic of pulse-labeling DNA strands. Bottom: DNA fibers were from RPE1 cells transfected with siRNAs and/or treated with B02 or vehicle as indicated. Red or Green portions of dual-labeled fibers were quantified and plotted as shown. Outliers were filtered using ROUT in Prism, ordinary one-way ANOVA followed by Tukey’s. ns – not significant, *P < 0.05, **P < 0.01, ***P < 0.001, ****P < 0.0001 **I.** Comet assay of hTERT-RPE1 cells treated with 2 mM HU for 40 h. Representative comets shown at the top. Indicated siRNAs were transfected two days before HU treatment. Red line represents median, n = 3 independent experiments, representative experiment shown. Mann Whitney test, ****P < 0.0001.

### RAD51 activity globally regulates stalled fork metabolism

RAD51 has been implicated in various fork processing roles in cells undergoing replication stress. It is important for fork reversal, protecting stalled forks against the activity of nucleases, and is key for recombination through template switching^51^. RAD51’s recombination function can be genetically uncoupled from the other functions, as multiple studies have shown that RAD51 strand exchange activity is not required for fork protection^50,52,53^. To determine if and how RAD51 regulates the phenotypes seen in the absence of SCAI/REV3, we first depleted RAD51 using a pool of siRNAs (Figure S6A). Depletion of RAD51 abolished the increased pRPA and γH2AX seen in the absence of SCAI (Figure S6A). We next tested RAD51’s role using two independent siRNAs and observe similar effects – loss of RAD51 abolished pRPA and γH2AX upon HU treatment even when SCAI was absent (Figure 6D). To further determine how RAD51 contributes to the increased pRPA and γH2AX seen upon fork stalling, we tested the effect of inhibiting RAD51 strand exchange activity using a previously described inhibitor B02^54^. Surprisingly, RAD51 inhibition completely abolished the break signaling seen upon loss of SCAI or REV3 (Figure 6E, compare lanes 2&3 to lanes 5&6), without affecting cell cycle distribution (Figure S6B), suggesting that this function requires or occurs downstream of the strand exchange activity of RAD51. We showed earlier that inhibition of fork reversal led to additive increases in fork breakage upon SCAI loss. We were curious whether this increase was also modulated by RAD51. Remarkably, RAD51 inhibition completely prevented the increased break signaling seen in the absence of SCAI when fork reversal was prevented either via SMARCAL1 knockdown (Figure 6F) or via PARP inhibition (Figure S6C) which has been shown to modulate fork reversal outcomes^15^.

Next, to determine how SMARCAL1 and SCAI modulated fork dynamics, we performed similar DNA fiber assays as in Figure 3A. SMARCAL1 loss has been shown to impair stalled-fork restart and promote replication stress associated fork breakage^55,56^. Depletion of SCAI and SMARCAL1 led to additively enhanced shortening of DNA fibers upon replication stress (Figure 6G, left panel). Co-depletion also led to reduced lengths of restarted forks (Figure 6G, right panel), suggesting that both proteins function in parallel to promote stalled fork rescue and prevent DNA breakage. Next, we tested the effect of RAD51 in a similar assay. Multiple groups have shown that RAD51 promotes restart of stalled forks, including through strand exchange-dependent mechanisms^13,52,57^. Unlike SMARCAL1, RAD51 inhibition rescued the fiber length shortening seen upon SCAI loss (Figure 6H, left panel) but markedly enhanced the shortening of the restarted forks (Figure 6H, right panel) suggesting that RAD51 promotes breakage of stalled forks but is essential for stalled fork rescue. To directly test RAD51’s effect on DNA break formation, we performed neutral comet assays. Upon replication stress, RAD51 depletion prevented DNA break formation in the absence of SCAI (Figure 6I, see also Figure S7B). Taken together these data suggests that RAD51 is important for formation of DNA breaks seen in the absence of Protexin, and that failure of fork reversal leads to increased RAD51 driven breakage when SCAI or REV3 is absent.

### BRCA1-RAD51 axis governs SLX1-SLX4 and ERCC1 mediated DNA breakage upon Protexin loss

We next wanted to determine how DNA breaks form in the absence of SCAI or REV3. Cells possess several structure specific nucleases that orchestrate cleavage of stressed forks^17^. Among these, MUS81 has been shown to promote fork processing and restart under various conditions^58,59^. Other nuclease complexes such as SLX4-SLX1 and XPF-ERCC1 have also been implicated ^17,60^. We co-depleted SCAI or REV3 cells with siRNAs against MUS81 or SLX4 following treatment with HU. We found that loss of SLX4, but not MUS81, reduced γH2AX levels in SCAI depleted cells (Figure 7A). A similar trend was observed in REV3 depleted cells: MUS81 had no effect or led to moderate increases in γH2AX, whereas SLX4 depletion reduced γH2AX levels (Figure 7B). We next determined whether SLX4-associated nuclease factors, specifically SLX1 or ERCC1, the binding partner of the XPF1 nuclease, also contributed to the elevated damage signaling caused by Protexin loss. We found reduced signaling when ERCC1 was knocked down and a similar but milder effect in the absence of SLX1 (Figure 7C). Having consistently observed involvement of SLX4, we directly examined break formation by performing neutral comet assays. As expected, SCAI depletion led to increased comet tail moments which was significantly reduced by SLX4 depletion (Figure 7D). Synchronization experiments revealed that this cleavage event occurred prior to mitotic entry (Figure S7B).

**Figure 7:**
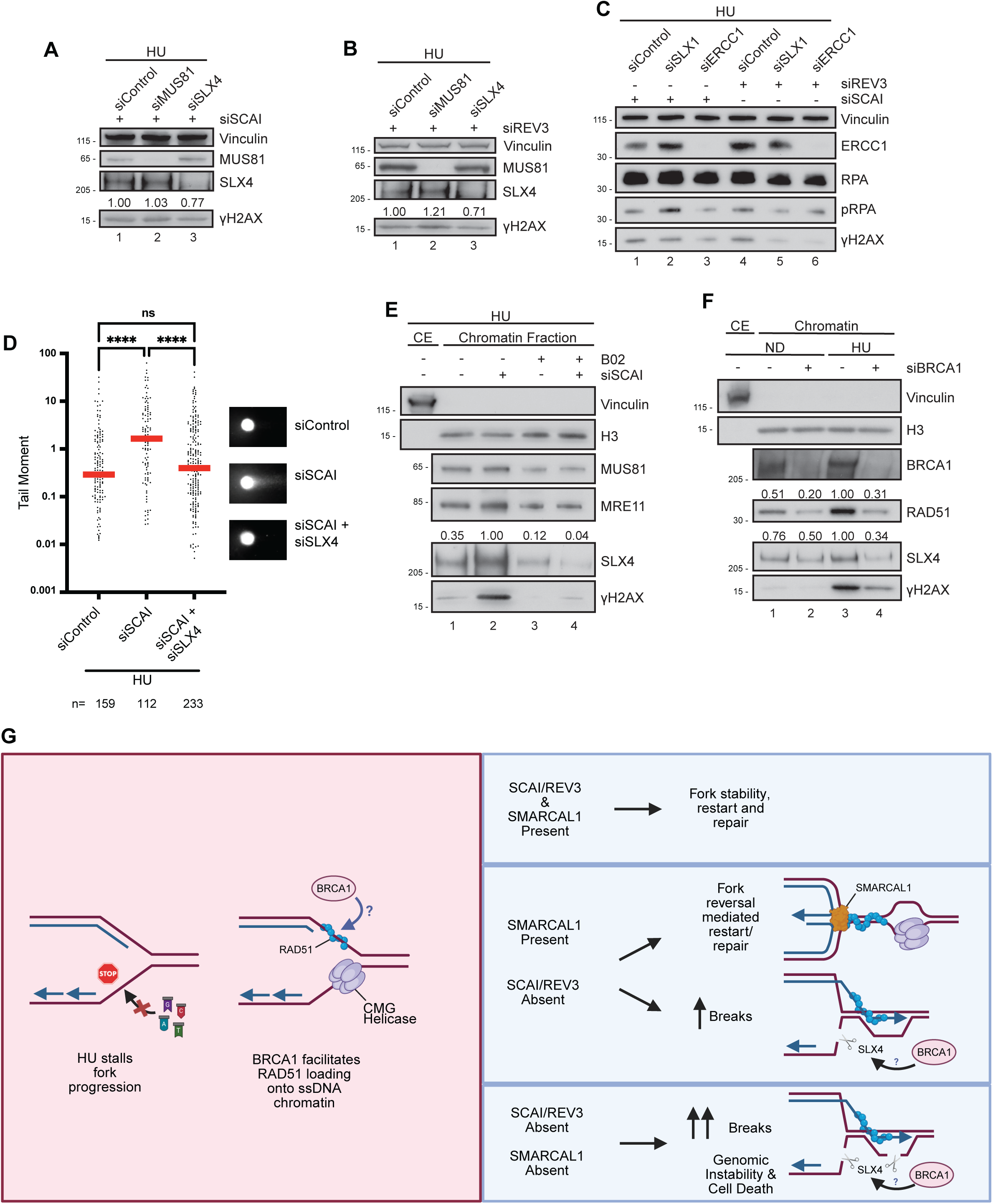
BRCA1-RAD51 axis regulate SLX1-SLX4 and ERCC1 mediated DNA breakage upon Protexin loss. **A.** WT RPE1 cells were transfected and treated with 2 mM HU for 18 h. WCLs were prepared, and WB were performed against the indicated proteins, n = 3 independent experiments. **B.** Western blots from WT RPE1 cells transfected with the indicated siRNAs and then treated with 1 mM HU for 18 h, n = 3 independent experiments. **C.** Western blots against the indicated proteins from WT RPE1 cells transfected, then treated with 1.5 mM HU for 18 h, n = 3 independent experiments. **D.** Comet assay of hTERT-RPE1 cells treated with 2 mM HU for 40 h. Indicated siRNAs were transfected two days before HU treatment. Representative comets are shown on the right. Red line represents median, n = 3 independent experiments, representative experiment shown. Kruskal-Wallis followed by Dunn’s multiple comparison test. ns – not significant. **E.** WT RPE1 cells were transfected with the indicated siRNAs, then treated with 1.5 mM HU for 18 h and fractioned prior to blotting for the indicated proteins. CE – cytoplasmic extract. SLX4 values are normalized to H3 loading control, n = 3 independent experiments. **F.** WT RPE1 cells were transfected and then treated with 2 mM HU for 18 h. Samples were processed for chromatin fraction prior to blotting for the indicated proteins. RAD51 and SLX4 values are normalized to H3 loading control, n = 3 independent experiments. **G.** Working model showing proposed roles of BRCA1, RAD51 and SCAI/REV3 upon fork stalling. See Discussion. Image was partially generated using BioRender.

We next sought to determine whether and how SLX4 was recruited to stalled forks and how this was modulated by loss of SCAI. Fractionation experiments revealed that SCAI loss led to increased amounts of SLX4 in the chromatin fraction compared to controls (Figure 7E, compare lanes 1 & 2). We did not observe such increases with MUS81 or MRE11 (Figure 7E). Strikingly, RAD51 inhibition led to significant reduction in not only γH2AX levels, but also in the amount of SLX4 on chromatin (Figure 7E), and the increased SLX4 amounts on chromatin seen in the absence of SCAI was prevented by B02 treatment (Figure 7E, compare lanes 2 and 4). Taken together, these experiments suggest that DNA breakage in the absence of SCAI/REV3 occurs downstream of RAD51 strand exchange activity but likely independent of fork reversal (see Figure 6).

To further characterize the mechanism by which stalled forks break, we depleted BRCA1. Upon SCAI depletion, BRCA1 KO showed reduced chromatin levels of SLX4 compared to controls (Figure S7A); this finding is consistent with a prior report ^61^. To further examine this role, and to determine how it was modulated by RAD51, we knocked down BRCA1 using siRNAs. HU treatment resulted in slightly increased amounts of SLX4 on chromatin (Figure 7E, compare lanes 1 & 3). Similar to what was seen following RAD51 inhibition or in BRCA1 KO cells, BRCA1 KD also resulted in reduction in the amount of damage induced SLX4 on chromatin (Figure 7F, compare lanes 1 & 3). Similar to what was seen following RAD51 inhibition or in BRCA1 KO cells, BRCA1 KD also resulted in reduction in the amount of damage induced SLX4 on chromatin (Figure 7F). Next, to determine how BRCA1 and RAD51 activities contributed to DNA breakage, we examined chromatin association of RAD51 and its modulation by BRCA1. Upon treatment with HU, there was increased amounts of RAD51 on chromatin (Figure 7F, compare lanes 1 & 3). Loss of BRCA1 significantly prevented the increase in RAD51 amounts on chromatin (Figure 7F, lane 4). Taken together these results identify a BRCA1-RAD51 axis upon Protexin loss that promotes SLX4 accumulation to salvage stalled replication forks.

## DISCUSSION

In this work, we have uncovered a novel stalled fork maintenance mechanism requiring the activities of both the fork protecting factor SCAI and the DNA polymerase REV3 to prevent pathologic fork breakage upon replication stress. Our experiments reveal key roles for BRCA1 and RAD51, as their loss prevented SLX4 chromatin association and DNA breakage in the absence of Protexin, suggesting failure downstream of HR. Supporting this, we find requirements for both the HR-promoting coiled-coil domain of BRCA1 as well as the strand exchange activity of RAD51. We propose that, at certain distressed forks, BRCA1-RAD51 drive fork restart that confers a dependency on the SCAI/REV3 complex. Surprisingly, this process appears to occur either parallel to or upstream of SMARCAL1/ZRANB3 or FBH1-mediated fork reversal, as loss of fork reversal factors led to increased break signaling, additive fork shortening and enhanced restart defects upon SCAI loss (model in Figure 7G).

Using multiple approaches, we show that increased break formation upon Protexin loss likely occurs in S phase, supporting the conclusion that Protexin limits fork breakage. Furthermore, single molecule fiber analysis revealed fork shortening upon HU treatment whenever SCAI or REV3 was absent. SCAI is known to restrict EXO1 activity, yet EXO1 depletion did not rescue fork shortening seen in the absence of SCAI under these conditions. In fact, loss of SMARCAL1, which could promote degradation of both nascent strands following reversal^35,49^, led to further reduction in fork lengths. The observation of reduced lengths of restarted forks alongside modest reduction in restart efficiency in the absence of SCAI suggest that distressed forks upon Protexin loss are salvaged for inefficient restart. This is likely due to RAD51 activity^13,52^ and suggests that SCAI may also function downstream of strand invasion. Indeed, restart experiments upon RAD51 inhibition show severely compromised fork restart efficiency whilst rescuing fork length shortening seen prior to release from replication stress.

In addition to its canonical roles in regulating resection and consequent pathway choice at conventional DSBs, BRCA1 has several proposed roles at stalled forks^62^. Its fork protection functions are well established, occurring downstream of fork reversal. Work in the literature has suggested roles for BRCA1 in stalled fork remodeling leading to fork restart^11,63,64^ with alternative mechanisms rescuing forks upon BRCA1 loss^65^. Our experiments allow us to uncouple BRCA1’s role from a fork protection function not only because BRCA1 loss paradoxically prevents DNA breakage but because depletion of fork reversal factors, which are necessary to provide a substrate for fork protection, does not abolish the DNA break formation seen in the absence of SCAI but rather increases it. We also show that this role occurs independently of or distinct from BRCA1’s known roles at conventional breaks, the C-terminus of BRCA1 is dispensable for this function, and we directly rule out a role for BRCA1’s resection partner, CtIP, in promoting break formation at stalled forks.

We previously employed a Tus/Ter reporter to show that SCAI is particularly important for HR promotion at stalled forks^22^. Our results here suggest that this role is important for fork maintenance and efficient fork restart, and that this role occurs downstream of RAD51 activity. We hypothesize that certain regions of the genome are prone to stall the advancing replication fork and require SCAI and REV3 to direct fork restoration and repair. The nature of such regions is not yet clear. Curiously, we observed a requirement for ERCC1 in driving increased break signaling in the absence of Protexin. Such a role has been shown by others^60,66^ and could suggest structural blocks such as hairpins. Indeed, using a fragile site reporter we found that SCAI/REV3 loss results in increased breaks^22^, consistent with our current findings. While SCAI and REV3 appear to function in concert, they do not have overlapping roles, and loss of SCAI seemed to lead to a stronger restart defect than loss of REV3. Future work will uncover under what scenarios SCAI and REV3 function together and how REV3 is regulated differentially from its role in translesion synthesis.

A central role for RAD51 in promoting fork restart has been proposed by others^20^, and the fork restart defects seen in the absence of SCAI appeared to occur downstream of RAD51 activity. BRCA1 depletion did not affect SCAI recruitment to chromatin, but results in significant reduction in RAD51 loading on chromatin. Whether BRCA1’s role here lies solely in RAD51 regulation is not yet clear. One interpretation of our data is that the SCAI/REV3 mediated restart mechanism occurs in parallel with SMARCAL1 /ZRANB3 or FBH1 mediated fork reversal. This is supported by the finding that loss of fork reversal factors like SMARCAL1 paradoxically led to increased break signaling in the absence of SCAI, which could indicate that forks normally processed via fork reversal are funneled into a SCAI dependent pathway that ultimately leads to collapse whenever SCAI was absent. Both pathways appear to require RAD51 activity, as inhibiting RAD51 prevented DNA breaks whenever SCAI and SMARCAL1 were co-depleted. RAD51 has been proposed to cause a CMG helicase-containing bubble ahead of annealed parental strands^57^. RAD51 could promote restart by invading newly synthesized DNA ahead of the annealed fork. Under-replicated regions can then be duplicated by gap-synthesis post repair. BRCA1 is not essential for fork reversal ^49^, but may enhance RAD51 chromatin association to drive strand invasion mediated fork restart. We show that SCAI is required for fork restart in this context, and in the absence of SCAI, SLX4 accumulates on chromatin. One possibility is that SLX4-SLX1 and ERCC1 promote cleavage of the resulting HR intermediates leading to collapse. Such broken forks can be engaged by alternative means of end joining or may persist leading to genomic instability and cell death. Whether the same RAD51 activity drives both reversal as well as HR-mediated fork restart will be a focus of future work.

## METHODS

### Cell lines

U2OS cells were originally obtained from ATCC (HTB-96). U2OS ΔSCAI and U2OS cells expressing lentiCRISPR control gRNAs (WT) were previously described^22^, and were passaged in McCoy’s 5A medium + L-Glutamine (16600-082, Gibco). hTERT-RPE1 cells were originally obtained from ATCC (CRL-4000). All RPE1 cell lines and their derivatives were passaged in DMEM/F12 (1:1) medium + L-Glutamine + 15mM HEPES (11330-032, Gibco). RPE1 ΔBRCA1/Δp53 (referred to as ΔBRCA1 in Figure 4) was a kind gift from Dr. Alan D’Andrea^67^. RPE1 TP53^-/-^ GFP PINDUCER20 (referred to as “parent” in Figures S5B and S7A), RPE1 TP53^-/-^ BRCA1^-/-^ GFP PINDUCER20 (referred to as “ΔBRCA1” in Figure 5C, 5D, S5B, and S7A), RPE1 Tp53^-/-^ BRCA1^-/-^GFP-BRCA1 PINDUCER20 (referred to as “WT” in Figures 5C, 5D, 5E and “GFP-BRCA1 Rescue” in Figure S5B) and RPE1 TP53^-/-^ BRCA1^-/-^ GFP-BRCA1 ΔCC PINDUCER20 (referred to as ΔCC) were kind gifts from Dr. Andrew Blackford and were previously described^48^.

### Reagents and Drugs

Dimethyl Sulfoxide (DMSO) was from Thomas Scientific (D0231). Hydroxyurea (HU) was obtained from Sigma-Aldrich (H8627-5G). Aphidicolin (APH) was from Sigma-Aldrich (A4487-1ML). The TLS inhibitor JH-RE-06 was obtained from MedChemExpress (HY-126214, 10mM/1mLSolution). Cisplatin was from Selleck Chemicals (S1166-50mg). Rad51 inhibitor B02 was from Sigma-Aldrich (553525-25MG). Olaparib from Selleck Chemicals (S1060-25mg). Colcemid was obtained from Millipore Sigma (10295892001).

### Antibodies

The antibodies used in this work include the following: Anti-vinculin 1:1000 (Sigma-Aldrich, V9131-.2ML), vinculin (7F9) 1:200 (Santa Cruz Biotechnology, sc-73614), SCAI 1:500 (Abcam, ab124688), Chk1 (2G1D5) 1:1000 (Cell Signaling, 2360S), P-Chk1 (S345) (133D3) Rabbit mAb 1:1000 (Cell Signaling, 2348S), RPA 32 kDa subunit (9H8) 1:200 (Santa Cruz Biotechnology, sc-56770, only used for western blots), RPA32 1:200 (Genetex, GTX70258, only used for IF), Rabbit x-Phospho RPA32 (S4/S8) 1:1000 (Bethyl, A300-245A), H2AX 1:1000 (Bethyl, A300-082A), Anti-phospho-Histone H2A.X (Ser139) 1:1000 (EMD Millipore Corp., 05-636), REV7 1:200 (Santa Cruz Biotechnology, sc-135977), Anti-Exonuclease 1 (EXO1) 1:1000 (Bethyl, A302-640A), ORC2 1:1000 (Abcam, ab68348), BRCA1 1:1000 (EMD Millipore Corp., 07-434), 53BP1 1:1000 (Bethyl, A300-272A), RIF1 1:1000 (Bethyl, A300-569A), CtIP 1:1000 (Bethyl, A300-488A), SmarcAL1 (A-2) 1:200 (Santa Cruz Biotechnology, sc-376377), ZRANB3 1:1000 (Bethyl, A303-033A), FBH1 1:1000 (Abcam, ab58881), RAD51 1:1000 (Abcam, ab63801), MUS81 1:200 (Santa Cruz Biotechnology, sc-53382), SLX4 1:1000 (Bethyl, A302-270A), H3 1:1000 (Cell Signaling, 9715), MRE11 1:1000 (Abcam, ab214), Goat anti-Mouse IgG (H+L) Secondary Antibody, HRP 1:2500 (Invitrogen, 31430), Goat anti-Rabbit IgG (H+L) Secondary Antibody, HRP 1:2500 (Invitrogen, 31460), Goat anti-Rabbit IgG (H+L) Highly Cross-Adsorbed Secondary Antibody, Alexa Fluor Plus 488 1:300 (Invitrogen, A32731), Goat anti-Mouse IgG (H+L) Highly Cross-Adsorbed Secondary Antibody, Alexa Fluor Plus 594 1:300 (Invitrogen, A32742), Goat anti-Mouse IgG (H+L) Highly Cross-Adsorbed Secondary Antibody, Alexa Fluor Plus 488 1:300 (Invitrogen, A32723).

### siRNA transfections

RPE1 and U2OS cells were seeded onto 6-well plates and reverse transfected with 20-40nM siRNA using Lipofectamine RNAiMAX reagent (13-778-150, Invitrogen) according to the manufacturer’s instructions. Cells were treated 48 h after treatment with drugs as indicated in the legend and processed 16 to 18 h later.

### SCAI rescue experiment

U2OS cells were transfected with plasmids using PolyJet reagent (SignaGen, SL10068B) according to the manufacturer’s instructions and treated with drugs as indicated in the Figure legends. SCAI was cloned into the pcDNA3.1 backbone as previously described ^22^. The ORF sequence of SCAI was modified to avoid being identified by gRNA in ΔSCAI cells. Specifically, GCTGAAGATGAT (Ala-Glu-Asp-Asp) was replaced with GCAGAGGACGAC using a site-directed mutagenesis kit (New England Biolabs, E0552S).

### Western blotting

Cells were harvested, washed with ice cold phosphate-buffered saline (PBS), and lysed in SDS lysis buffer (50 mM Tris HCl pH 6.8, 100 mM NaCl, 1.7% SDS, 10 mM NaF, 7% glycerol) prior to sonication. Protein content was measured using Bradford reagent (23236, Life Technologies) on an Eppendorf Biophotometer. Laemmli sample buffer (1610747, Bio-Rad Laboratories) was added with 20 mM DTT prior to boiling the samples for 5-10 min. Whole cell lysates were analyzed using sodium dodecyl sulfate gel electrophoresis (SDS-PAGE) and transferred onto nitrocellulose membranes. Membranes were blocked with 5% (wt/vol) milk in Tris-buffered saline with Tween (TBST) for a minimum of 30 min and then probed with the indicated antibodies. Protein bands were quantified with ImageJ.

### Chromatin fractionation

Chromatin fractions were done as previously described ^68^. Briefly, RPE1 cells previously transfected with the indicated siRNAs were seeded into 10 cm plates, treated with drugs as indicated, then harvested and washed with ice cold PBS. Sedimented cells were resuspended in cold Solution 1 (10 mM Hepes (pH 7.9), 10 mM KCl, 1.5 mM MgCl_2_, 0.34 M sucrose, 1 mM DTT, 10% glycerol, 1mM Na_3_VO_4_, protease and phosphatase inhibitors). Triton X-100 was added to 0.1%, and samples were incubated on ice prior to centrifuging. The supernatant was removed as the cytoplasmic extract, and sedimented nuclei were washed with Solution 1 prior to being lysed in Solution 2 (3 mM EDTA, 0.2 mM EGTA, 1 mM DTT, protease and phosphatase inhibitors) on ice. Samples were centrifuged, washed with cold Solution 2, and then the chromatin-enriched pellets were lysed with SDS lysis buffer. Chromatin extract samples were sonicated, and protein content was measured. Sample buffer was added to all samples and boiled prior to western blotting.

### Cell cycle analyses

Cells were treated with indicated drugs for 18h prior to being harvested. Cells were then washed with ice cold PBS, then fixed in 70% ethanol at 4 °C overnight. Samples were pelleted, washed with ice cold PBS, and resuspended in 50 µg/mL propidium iodide solution containing 0.1 mg/mL RNase A. Samples were then incubated at 37°C in the dark for 15-30 min and flow cytometry was performed using a FACSymphony A5 (BD Biosciences). Data were analyzed using FlowJo software.

### Immunofluorescent staining for Flow Cytometry

RPE1 cells were reverse transfected with the indicated siRNAs as described. 48 or 64 hours post transfection; cells were left untreated (0 hr control) or treated with 1.5 mM HU for the indicated duration. After the treatment duration, cells were harvested, fixed and immuno-labelled following the protocol described by Zimmermann *et al* with minor modifications. Briefly, cells were detached using trypsin, pelleted at 600 rpm for 5 min, washed in 1X PBS then fixed in 1 ml of 4% PFA for 10 min at RT. The cells were again pelleted as before and resuspended in 100 µl of PBS. 900 µl of prechilled (-20oC) methanol was then added to the cell suspension dropwise while vortexing at the lowest speed. Fixed cells were then kept at -20oC at least overnight. Next, cells were spun down, washed in PBS, resuspended in blocking buffer (3% BSA in PBS) and incubated for 30 minutes at room temperature. After blocking, cells were pelleted and resuspended in 200 µl of dilute rabbit anti-pRPA antibody (Bethyl A300-245A; 1:750) and mouse anti Phospho-H2A.X(γH2A.X) (EMD Millipore 05636IMI 1:750) in blocking buffer and incubated at RT for 2 hrs. Following the 2-hour incubation, the antibody was diluted by adding 2 ml of PBS and cells spun down. The supernatant was carefully removed, and the pellet resuspended in 200 µl of dilute Alexa Fluor 488-conjugated goat anti-rabbit antibody and Alexa Fluor594-conjugated anti-mouse (Molecular Probes/Thermo Fisher, 1:1,000) in blocking buffer then incubated at RT for 1 hour. The cells were washed by adding 2 ml of PBS and centrifuging for 5 minutes. The pellet was resuspended in 1 µg/ml DAPI incubated for 15 minutes and samples analyzed on the FACSymphony A3 (BD Biosciences). Data was acquired using the BD FACSDiva software and analyzed using FlowJo.

### Clonogenic survival assays

Cells were transfected as stated above for 48 h with the indicated siRNAs and then treated with indicated doses of drugs for 18 h. The number of cells plated in six-well plates was adjusted based on the dose of drug the cells were exposed to. Cells were allowed to form colonies for 10-14 days prior to being fixed and stained using Crystal Violet. Analyses were performed using ImageJ.

### Immunofluorescence Assay

Cells were transfected with the indicated siRNAs as above and plated onto glass coverslips 18 h later. 48 h after the transfection, the cells were treated with HU for the indicated dose and duration. The cells were then washed with ice-cold PBS on ice, pre-extracted for 5 min on ice using cytoskeleton buffer (100 mM NaCl, 10 mM HEPES (pH 7.8), 3 mM MgCl_2_, 300 mM sucrose, 0.1% Triton X-100), fixed with 4% paraformaldehyde in PBS for 10 min at room temp and then permeabilized with 0.5% Triton-100 in 1X PBS. Samples were then blocked with 3% bovine serum albumin (BSA) in PBS for a minimum of 30 min on a rocker followed by overnight incubation with primary antibodies at 4 °C in a humidified chamber. Unbound primary antibodies were washed using 3% BSA in PBS and then samples incubated with secondary antibody for 1 h at RT in the dark. Unbound antibody was washed with PBS and then samples stained with 1 μg/ml DAPI for 1 h before mounting the coverslips on a microscope slide for imaging. Images were acquired using an LSM 780 NLO (Zeiss), and a minimum of 100 nuclei for each group were analyzed using ImageJ.

### Proximity Ligation Assay

RPE cells were reverse transfected with the indicated siRNAs. 24h later cells were replated on coverslips and treated with drugs as indicated. Coverslips were washed once with 1xPBS, and pre-extracted with CSK (100nM NaCl, 100mM HEPES pH 7.8, 3mM MgCl2, 300mM Sucrose, 0.1% TritonX-100) for 5min on ice and then with CSK+0.5% TritonX-100 for 5min on ice. Samples were fixed with 100% Methanol for 20min at -20°C. Coverslips blocked with 5% BSA at room temperature for at least 1 hr. Indicated primary antibodies were incubated overnight at 4°C in a humid chamber, washed with 1xPBS. Subsequent steps were performed according to the manufacturer’s (Duolink®, Sigma) instructions. Briefly, cells were incubated with secondary PLA probes (Duolink® In Situ PLA® Probe Anti-Rabbit PLUS (DUO92002), Duolink® In Situ PLA® Probe Anti-Mouse MINUS (DUO92004) in antibody diluent) for 1h at 37°C in a humid chamber, washed twice with Wash Buffer A (10mM Tris Base, 150mM NaCl, 0.05% Tween-20, pH=7.4) and then incubated with Duolink® In Situ Detection Reagents Red, first with the ligation mix for 30min at 37°C in a humid chamber. Cells were then washed twice with 1mL Wash Buffer A and incubated with PLA amplification for 100min at 37°C in a humid chamber. Coverslips were washed with Wash Buffer B (200mM Tris Base, 100mM NaCl, pH 7.4) and then 0.01x Wash Buffer B before being sealed with DAPI in mounting media (SouthernBiotech, DAPI Fluoromont-G, 0100-20). PLA foci were imaged with a 40x oil objective on a Zeiss 780 confocal microscope. Images were taken as 10x0.52µm Z-stacks to ensure the entire nucleus was imaged, and projected using FIJI sum slices for intensity and count analysis. Representative images were max projections.

### QIBC image acquisition and analysis

Fixed and stained slides were imaged on a Molecular Devices ImageXpress Microscope equipped with a Nikon 40x/0.95 PlanApo objective. Around 2000 nuclei were acquired for each condition. Quantification was then performed using the “Multi Wavelength Cell Scoring” module in the MetaXpress software. Briefly, nuclei were segmented using the DAPI channel, and the raw Mean Fluorescent Intensities (MFIs) of each nucleus were measured for DAPI, RPA (AF488), and gamma-H2AX (AF594). All cells were then plotted for each condition and p-values generated by Ordinary one-way ANOVA, Tukey’s multiple comparison test

### DNA fiber assays

hTERT-RPE1 cells were reverse transfected with the indicated siRNAs at 20 nM using Lipofectamine RNAiMAX reagent (Invitrogen). Transfected hTERT-RPE1 cells were pulse-labeled with CldU followed by IdU as indicated in each Figure. After being labeled, cells were harvested and embedded in 0.6% low melting point agarose gel plugs. Cells in plugs were incubated with lysis solution (0.2 M EDTA, 0.5% SDS, 3 mg/mL Proteinase K) overnight at 50°C. Each plug was dissolved in 0.5 M MES solution (pH 5.5) by β-agarase (NEB) overnight treatment and DNA was attached to a pre-treated coverslip (PolyAn) using FiberComb® Molecular Combing System (GENOMIC VISION). DNA on coverslips was denatured by incubating in denaturing solution (0.5 N NaOH, 1 M NaCl) for 8 min. Blocking was performed with 5% BSA in PBS for 60 min. For immunostaining, coverslips were incubated overnight with primary antibodies; ab6326 anti-BrdU (cross-reacts with CldU) antibody (rat) (1:50) and BD Biosciences 347580 anti-BrdU (cross-reacts with ldU) antibody (mouse) (1:8). Slides were washed with PBS followed by incubation for one hour with the secondary antibodies; anti-rat AIexa-488 antibody (1:300) and anti-mouse Alexa-594 antibody (1:300). After wash with PBS, coverslips were mounted onto glass slides with anti-fade mounting medium (SlowFade™ Glass Soft-Set Antifade Mountant, Invitrogen). Images were acquired with Leica STELLARIS confocal microscope. DNA fiber numbers and lengths were manually analyzed using LAS X software (Leica Microsystems).

### Metaphase spreads

hTERT-RPE1 cells were reverse transfected with the indicated siRNAs as above using Lipofectamine RNAiMAX. Two days after the transfection, cells were treated with 2 mM HU or PBS for 18 h, followed by PBS wash and 24 h incubation in normal medium. Cells were exposed to colcemid (100 ng/ml) for the last 3 h of the 24 h normal medium incubation before harvest. Collected cells were treated with a hypotonic solution (1:2 mixture of 0.075 M KCl and 0.9% Na citrate) for 10 min and fixed with 3:1 methanol: acetic acid. Fixed cells were transferred on glass slides dropwise, and the slides were stained with Giemsa stain. Cells in metaphase were observed using a Zeiss Axiovert 200M microscope and captured using AxioVision. Over 50 metaphase cells were analyzed per sample.

### Neutral Comet assay

hTERT-RPE1 cells were incubated with 2 mM Thymidine for 16 h followed by reverse transfection with indicated siRNAs at 20 nM using Lipofectamine RNAiMAX reagent (Invitrogen). Eight hour after the end of the first Thymidine incubation, cells were incubated with 2mM Thymidine again for 16 h. HU 2 mM treatment started 2 h after the end of the second Thymidine incubation. Cells were harvested 24 h later for Comet assay. Alternatively, hTERT-RPE1 or U2OS cells were reverse transfected as above then, two days after the transfection, treated with 2 mM HU or PBS for 40 h. After HU treatment, cells were harvested and resuspended in ice-cold PBS. 10μL of the cell suspension was mixed with 100 μL of melted 0.7% low melting point agarose gel, and the mixture was spread onto a pre-coated CometSlide™ (R&D SYSTEMS®). After the agarose gel was solidified at 4 °C, the slide was immersed in lysis solution (10 mM Tris base, 2.5 M NaCl, 100 mM Na_2_-EDTA, 10% DMSO, 1% Triton X-100, pH 10) for 2 h at 4 °C followed by a 30 min incubation in neutral electrophoresis buffer (100 mM Tris base, 300 mM sodium acetate, pH 9). Then, the slide was placed in an electric field for 40 min. After the electrophoresis, the slide was immersed in DNA precipitation solution (82% ethanol, 1 mM ammonium acetate) for 30 min followed by a 30 min incubation in 70% ethanol. SYBR Gold dye (Invitrogen) was used to stain DNA (“comet”) and images were obtained with TissueFAXS (TissueGnostics). Comets were analyzed with ImageJ (OpenComet).

### Quantification and statistical analyses

Statistical analyses were performed using Prism 10 (GraphPad). All statistics used are indicated in the Figure legends. For all experiments: n.s. P ≥ 0.05; *P<0.05; **P < 0.01; ***P < 0.001; ****P<0.0001. All experiments in this work were performed multiple independent times with similar results. Multiple siRNAs, multiple knockout clones, and multiple cell lines were analyzed to confirm that results were not caused by off-target effects or clonal variations. Representative images and/or experiments are shown for westerns, cell cycle analyses, DNA fiber assays and IFs.

## Acknowledgements

We thank all the members of the Adeyemi laboratory for advice and helpful discussions. We thank Dr. Blackford and Dr. Setiaputra for providing reagents.

## Author Contributions

Conceptualization: R.O.A., Methodology: C.U., H.T., R.B., S.K., X.J., R.O.A., Investigation: C.U., H.T., R.B., S.K., X.J., R.O.A., Visualization: C.U., H.T., R.B., S.K., X.J., R.O.A., Supervision: R.O.A., Writing—original draft: R.O.A., Writing—review & editing: C.U., H.T., R.B., S.K., X.J., R.O.A.

## Data and Materials Availability

All data needed to evaluate the conclusions in the paper are present in the paper and/or the Supplementary Materials. Materials resulting from the findings of this study are available from the corresponding author upon request.

Figure S1 A. WT RPE1 cells were transfected, then treated with the indicated doses of HU for 18 h and allowed to form colonies. Mean ± SD shown, n = 2 independent experiments. **B.** Representative images from (**A**) are shown. **C.** Immunofluorescence (IF) analyses showing representative RPA foci in WT RPE1 cells post-treatment with indicated siRNAs. Cells were treated with 1 mM HU for 18 h prior to IF. Merge includes RPA foci and DAPI. Scale bars indicate 5 micron. Intensity of nuclear RPA staining was quantified using ImageJ and plotted on the right. ****P < 0.0001 and **P = 0.0063, Ordinary one-way ANOVA followed by Tukey’s. **D.** Representative IF images showing γH2AX foci in WT RPE1 cells following treatment with the indicated siRNAs. Cells were treated with 1.5 mM HU for 18 h prior to IF analyses. Scale bars indicate 5 micron. Quantification shown on the right. ****P < 0.0001, ANOVA followed by Tukey’s. **E.** Western blots against the indicated proteins from WT RPE1 cells pretreated with 10 μM TLS inhibitor JH-RE-06 or vehicle (DMSO) for 30 min, then treated with 1.5 mM HU for 18 h, n = 3 independent experiments. **F.** WT and SCAI-null U2OS cells were treated with the indicated siRNAs, then treated with 1 mM HU for 18 h prior to blotting for the indicated proteins, n = 3 independent experiments.

Figure S2 A. Representative images of 1.5 mM HU-treated RPE1 cells with control or siSCAI at the indicated timepoints showing γH2AX, PCNA, and DAPI staining. Scale bars indicate 50 micron. Note that representative images at the 8 h timepoint is included in the main text with the same images. **B.** Parallel western blot showing SCAI knockdown in RPE1 cells harvested at 8 h after HU. **C.** Quantification of γH2AX intensity from (**A**). T-test, *P < 0.05. Black bars represent mean. **D.** Pearson’s correlation coefficient between γH2AX and PCNA measured using BIOP JACoP FIJI plugin using manual thresholding based on control images (γH2AX = 36822; PCNA = 26354). Pearson’s correlation was recorded as a single point for each image (n = 14-16) with 10-16 cells per image. Red bars represent mean. **E.** Plot showing the observed overlap between γH2AX and PCNA staining from (**A**), thresholded for both channels at 1.25x10^8 a.u. (dotted line shown in (**C**)). Expected values were obtained by multiplying γH2AX and PCNA positive fractions. **F.** WT RPE1 cells were transfected with the indicated siRNAs and then treated with 2 mM HU for the indicated times. WB were performed against the indicated proteins, n = 3 independent experiments. **G.** Schematic of double thymidine block protocol for cell synchronization. WT RPE1 cells treated with 2 mM thymidine for 16 h, transfected and released for 8 h, treated with 2 mM thymidine for another 16 h, released for 2 h, then harvested for cell cycle analysis by flow cytometry. Representative images shown below. **H.** PLA foci between PCNA and γH2AX on siSCAI 16 h post treatment with 1.5 mM HU. **I.** Quantification of (**H**) with Mean ± SEM for n = 2 independent experiments, with >180 cells/condition/experiment. All experiments were analyzed with One-way ANOVA with a Kruskal-Wallis test for multiple comparisons, *P < 0.0332, **P < 0.0021, ***P < 0.0002, ****P < 0.0001. **J.** Schematic of cell treatment protocol to perform double thymidine block, transfect, and synchronize cells. Comet assay of WT RPE1 cells treated with 2 mM HU for 24 h or untreated. Red line represents median, n = 3 independent experiments, representative experiment shown. Kruskal-Wallis followed by Dunn’s multiple comparison test, ****P < 0.0001.

Figure S3 A. Schematic of pulse-labeling DNA strands. DNA fibers were from RPE1 cells transfected with siRNAs as indicated. New origin firing events (red only) on left panel, percentage of red labels over total number of fibers on the right. Mann-Whitney test, n = 3 independent experiments, ns = not significant. **B.** Schematic of pulse-labeling DNA strands and lengths of two-color DNA fibers. DNA fibers were from hTERT-RPE1 cells transfected with siRNAs as indicated. Red line represents median, n = 2 independent experiments, representative experiment shown. Kruskal-Wallis followed by Dunn’s multiple comparison test. *P < 0.05, ****P < 0.0001. **C.** WT RPE1 cells were transfected with the indicated siRNAs, then treated with 1.5 mM HU for 18 h and assayed for the indicated proteins by WB, n = 3 independent experiments.

Figure S4 A. WT RPE1 cells were transfected with the indicated siRNAs, then treated with 2 mM HU for 18 h before assaying for the indicated proteins by WB, n = 3 independent experiments. **B.** Western blots from WT and SCAI-null U2OS cell lines transfected with the indicated siRNAs and then treated with 1.5 μM cisplatin for 18 h, n = 3 independent experiments. **C.** WT and BRCA1-null RPE1 cells were transfected with the indicated siRNAs, then treated with 1 mM HU for 18 h and fractioned prior to blotting for the indicated proteins. SCAI values are normalized to ORC2 loading control, n = 3 independent experiments. **D.** Comet assay of WT RPE1 following synchronization and siRNA transfection as in **Figure S2J**. Cells treated with 2 mM HU for 24 h. Red line represents median. Kruskal-Wallis followed by Dunn’s multiple comparison test. ****P < 0.0001 **E.** Representative images showing cell cycle distribution in WT and BRCA1-null RPE1. Quantified on the right, mean ± SD shown, n = 2 independent experiments.

**Figure S5 A.** GFP-WT-BRCA1 rescue (labeled WT), BRCA1-null (Δ), and GFP-BRCA1 Rescued RPE1 cells were assayed for the indicated proteins by WB. **B.** P53-null RPE1 parent (Parent), BRCA1-null (ΔBRCA1), and GFP-WT-BRCA1 Rescue (GFP-BRCA1 Rescue) RPE1 cell lines were transfected and treated with 1 mM HU for 18 h. WCLs were prepared, and WB were performed against the indicated proteins, n = 2 independent experiments.

Figure S6 A. WT RPE1 cells were transfected with the indicated siRNAs, then treated with 1 mM HU for 18 h and assayed for the indicated proteins by WB, n = 3 independent experiments. **B.** Representative images showing cell cycle distribution in WT RPE1 cells 18 h following treatment with 27 μM B02 compared to vehicle controls. Quantified on the right, mean ± SD shown, n = 2 independent experiments. **C.** Western blots from WT RPE1 cells transfected with the indicated siRNAs, pretreated with DMSO, 10 μM Olaparib (Olap), or 27 μM B02, and then treated with 1 mM HU for 18h, n = 3 independent experiments.

**Figure S7 A.** Parent and BRCA1-null RPE1 cells were transfected and then treated with 3 mM HU for 18 h. Samples were processed for chromatin fraction prior to blotting for the indicated proteins, n = 2 independent experiments. **B.** Comet assay of WT RPE1 following synchronization and siRNA transfection as in **Figure S2J**. Cells treated with 2 mM HU for 24 h. Red line represents median. Kruskal-Wallis followed by Dunn’s multiple comparison test. ****P < 0.0001.

